# Classes for the masses: Systematic classification of unknowns using fragmentation spectra

**DOI:** 10.1101/2020.04.17.046672

**Authors:** Kai Dührkop, Louis Felix Nothias, Markus Fleischauer, Marcus Ludwig, Martin A. Hoffmann, Juho Rousu, Pieter C. Dorrestein, Sebastian Böcker

## Abstract

Metabolomics experiments can employ non-targeted tandem mass spectrometry to detect hundreds to thousands of molecules in a biological sample. Structural annotation of molecules is typically carried out by searching their fragmentation spectra in spectral libraries or, recently, in structure databases. Annotations are limited to structures present in the library or database employed, prohibiting a thorough utilization of the experimental data. We present a computational tool for systematic compound class annotation: CANOPUS uses a deep neural network to predict 1,270 compound classes from fragmentation spectra, and explicitly targets compounds where neither spectral nor structural reference data are available. CANOPUS even predicts classes for which no MS/MS training data are available. We demonstrate the broad utility of CANOPUS by investigating the effect of the microbial colonization in the digestive system in mice, and through analysis of the chemodiversity of different Euphorbia plants; both uniquely revealing biological insights at the compound class level.

## INTRODUCTION

Liquid chromatography mass spectrometry (LC-MS) can detect large numbers of small molecules from fractional amounts of samples, and has been widely adopted by the metabolomics community. LC-MS, when performed in an untargeted fashion, allows us to detect hundreds to thousands of metabolites from a single sample, but annotation of these metabolites remains highly challenging. Annotations are typically reached by searching fragmentation spectra (MS/MS) in spectral libraries^1, 2^ or, more recently, structure databases^3^. Yet, only a fraction of metabolites can be annotated in this fashion^4^. Spectral libraries are clearly limited in size and largely have been created with commercially available compounds ^5^; but even molecular structure databases, which can be orders of magnitude larger, usually do not cover all biomolecular structures from a particular sample. We do not expect that this situation is going to change fundamentally during the next decade. This means that for most of the data that is acquired in such metabolomics experiments, little structural insights can be obtained. In particular, it is currently not possible to obtain a comprehensive structural picture of which metabolites are present in a sample, and which are up- or down-regulated between two experimental conditions. Here, we present CANOPUS, a computational method that addresses this problem by assigning compound classes to every metabolite MS/MS feature in an LC-MS/MS run; this includes “unknown” metabolites with structures not recorded in any database or publication.

The problem of predicting the presence or absence of certain substructures has been considered since the 1960s, predominantly for GC-MS data^6-8^. FingerID^9^ and CSI:FingerID^10^ predict molecular fingerprints that encode several hundreds or thousands of substructures in a molecule, respectively. Substructures describe local parts of the molecule, such as the presence of a hydroxy group; in contrast, compound classes are usually substantially more complex. Compound classes have been defined in, say, the ChEBI ontology^11^ or the MeSH thesaurus^12^, but class annotations are available for only a small fraction of molecular structures. In contrast, ClassyFire^13^ enables deterministic assignment of classes solely from structure, allowing us to classify *all* molecular structures. ClassyFire definitions involve the use of logical expressions, substructures with variable length, and substructure count constraints.

To understand what compound classes are detected from a sample, we need a method that can predict these classes directly from MS/MS data. Unfortunately, this task comes with fundamental challenges which have never been addressed so far, and which make class assignment an even harder problem than searching in molecular structure databases: First, can we predict classes for which limited or no MS/MS data are available in spectral libraries? As libraries are notoriously incomplete and most classes are sparse, we have, for the majority of classes, minimal reference data belonging to the class. The second challenge is more subtle and simultaneously more problematic: Even if enough MS/MS data appear to be available for some class, it is possible that compounds for which we have reference spectra are not distributed evenly (or not at all) among subclasses. As an extreme case, assume our training data contain reference spectra for *pregnane steroids* but no other *steroids*. Training a model for *steroids* using this data will in fact only predict the subclass *pregnane steroids*, whereas all compounds from other subclasses (say, *ergostane steroids*) will be misclassified as “not a *steroid*” *with high confidence*. We will show that this seemingly theoretical issue can have massive practical consequences. For known class/subclass pair, we can identify and point out such “misguided classifiers” from the training data; but this is not possible for class/subclass pairs that are not part of the classification scheme. Third, can we predict classes for a compound that is truly unknown, meaning that not even its molecular structure has been recorded in any structure database? These “novel compounds” are arguably the ones for which class prediction is desired most; but how can we *evaluate* whether a method is able to target such “novel compounds”?

At present, three strategies for structural classification exist: a) Cluster compounds based on spectral similarity, then propagate compound class annotations from database search in a semi-automated manner^14-16^ b) Search for the query compound in a spectral library^17, 18^ or a structure database^19, 20^; consider the top *k* hits for assigning compound classes. c) Use machine learning methods to directly predict compound classes from the MS/MS spectrum^19, 21^. None of these strategies can address all challenges mentioned above, as we detail in the Methods section; furthermore, no ready-to-use computational tools for automated compound class annotation from LC-MS data are publicly available.

## Results

Here, we present the two-step approach CANOPUS (Class Assignment aNd Ontology Prediction Using mass Spectrometry) that addresses all of the abovementioned challenges. The workflow of CANOPUS is depicted in Fig. 1: Given an MS/MS spectrum as input, we use a series of support vector machines (SVM) to predict establish a probabilistic fingerprint of the query compound^9, 10^. This probabilistic fingerprint is used as input of a Deep Neural Network (DNN)^22^, which then predicts all compound classes simultaneously. SVMs are trained using reference MS/MS spectra; in contrast, the DNN is trained on 1.11 million compound structures and does not require any MS/MS data. To train the DNN, we have to simulate a “realistic” probabilistic fingerprint for any given molecular structure, although no MS/MS data for this structure is available. This integration of two machine learning techniques allows CANOPUS to reach high-quality predictions for 1,270 compound classes. Because the predictions are now independent from the availability of MS/MS reference data, CANOPUS can predict compound classes even when there are no MS/MS spectra for training the method. Equally important, it can predict classes for which MS/MS training data is missing for a complete subclass. Uniquely, CANOPUS permits a global over view of the compound classes measured in a biological sample, but also the differences between cohorts at the compound class level

**Figure 1.**
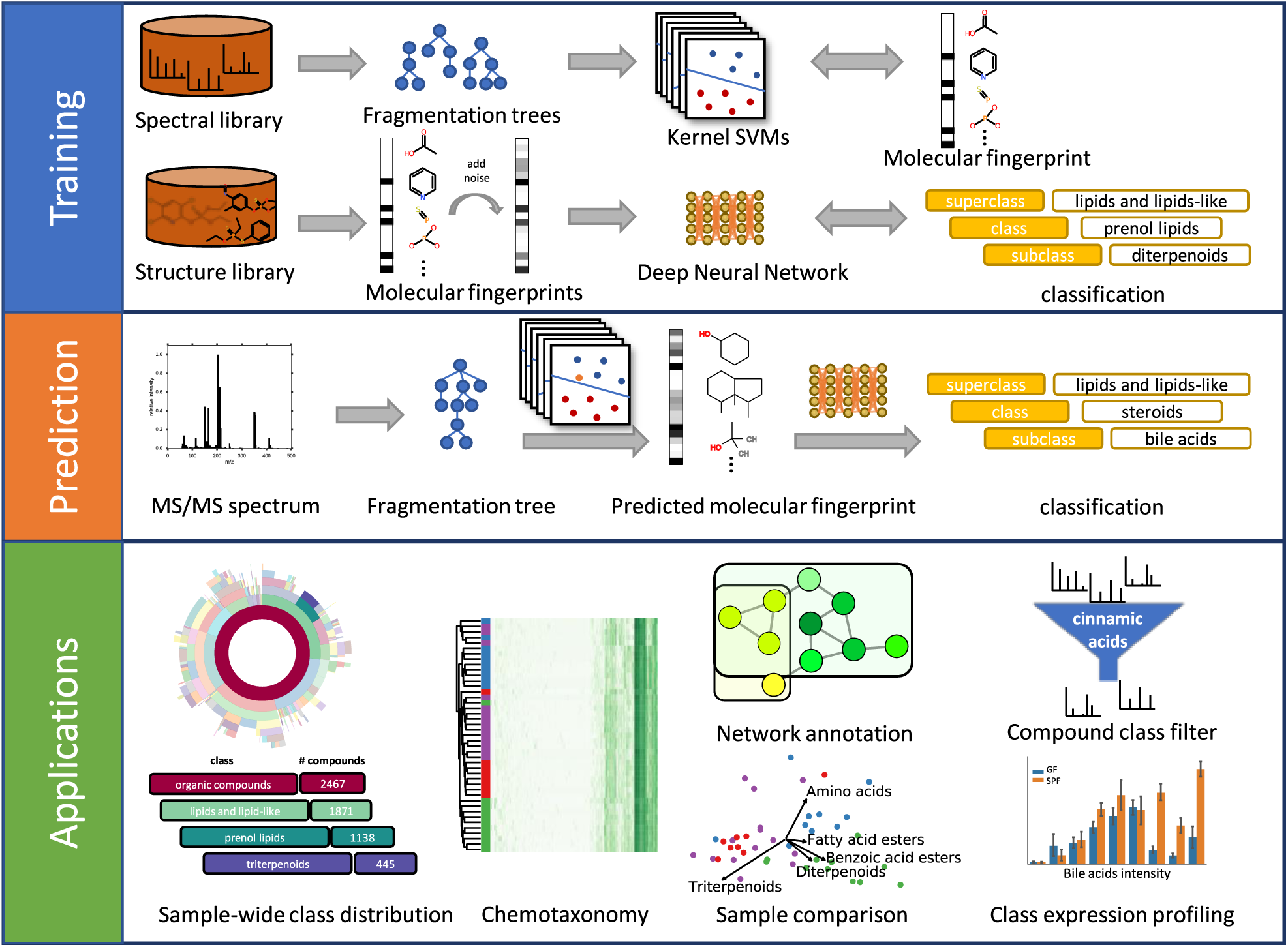
CANOPUS workflow. In the training phase, we train a battery of kernel support vector machines for predicting molecular fingerprints from fragmentation spectra, and we train one deep neural network to predict compound classes from molecular fingerprints (multi-label classification). In the prediction phase, we classify the query compound from its MS/MS spectrum by computing the fragmentation tree, predicting its molecular fingerprint, and predicting the compound classes from the fingerprint using the deep neural network. Numerous applications of compound classification exist, some of which are highlighted throughout this paper.

### CANOPUS evaluation

To evaluate CANOPUS we used reference MS/MS libraries, as we need to know the true answer for evaluation. The kernel SVMs for predicting the molecular fingerprint were trained on MS/MS spectra from 23,965 compounds. We used structure-disjoint ten-fold cross-validation on the training data, and structure-disjoint evaluation on an independent MS/MS dataset of 3,385 compounds. This ensures that all compounds are *novel* in evaluation, meaning that no reference MS/MS spectra are available for dereplication. The DNN was trained on 1.12 million structures with compound classes assigned by ClassyFire^13^. All structures for which MS/MS spectra are available, were removed from the DNN training data. Again, this ensures that all compounds are truly *novel* in evaluation, meaning that *both the MS/MS and the structure* are unknown to the method.

For compounds from the SVM training dataset, CANOPUS reached an average accuracy of 99.4%, and 1,094 of 1,270 classes were predicted with accuracy at least 99%. However, compound classes are very sparse, so the Matthews Correlation Coefficient (MCC) is a more suitable performance measure^23^. The MCC is between -1 and +1; it is 0 if a classifier performs no better than random, and +1 for a perfect classifier. See Supplementary Figure 1 for the MCC of all 732 compound classes with at least 50 examples. CANOPUS predicted 370 compound classes with MCC at least 0.8 (Fig. 2a, Supplementary Material 1), including *phosphocholines* (MCC 0.954), *flavonoid O-glycosides* (MCC 0.918), and *pregnane steroids* (MCC 0.867); 943 of 1,270 classes can be predicted with MCC at least 0.5.

**Figure 2.**
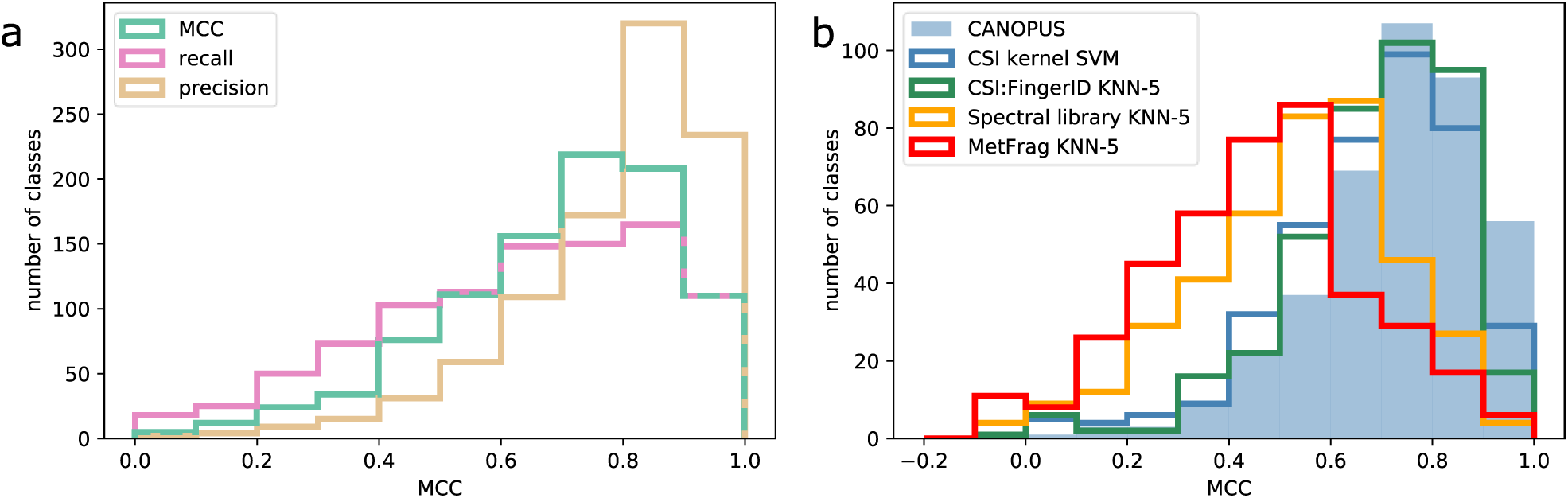
Method evaluation histogram plots. (a) Histogram of Matthews Correlation Coefficients (MCC), recall, and precision for CANOPUS; SVM training dataset, compound classes with at least 20 examples. (b) Histogram of MCC for four methods; independent dataset, compound classes with at least 20 positive examples. Recall that MCC of 0 corresponds to random predictions. “CSI Kernel SVM” is the direct prediction from MS/MS spectra using a kernel SVM. “MetFrag KNN-5” and “CSI:FingerID KNN-5” search in PubChem using MetFrag or CSI:FingerID, “Spectral library KNN-5” searches in the SVM training dataset using cosine similarity. All KNN-5 methods use majority vote of the top 5 search results for each compound class.

For the independent dataset, CANOPUS reached an average prediction accuracy of 99.4%, and 1,093 classes were predicted with accuracy at least 99% (Supplementary Material 2). The independent dataset is substantially smaller than the SVM training dataset; but MCC estimation can be inaccurate and misleading if too few positive examples are available. We therefore distinguished between *rich classes* with at least 20 positive examples (400 classes), and *sparse classes* with fewer examples (870 classes). The average MCC for rich classes was 0.729 (Fig. 2b). For the sparse classes, we computed the microaveraged MCC, which sums up true positives, true negatives, false positives, and false negatives over all sparse classes. The microaveraged MCC for all sparse classes was 0.531.

We further evaluated CANOPUS against four other methods for compound classification (Fig. 2b). The first two methods search in a structure database using an *in silico* tool, then perform a majority vote of the top 5 candidates for each compound class individually. We evaluated two widely used *in silico* tools, MetFrag^24, 25^ (MetFrag KNN-5) and CSI:FingerID^10, 26^ (CSI:FingerID KNN-5) for this purpose. These methods cannot provide any classifications if the query molecular formula is absent from the structure database. Third, we evaluated against searching in a spectral library (Spectral library KNN-5). We do not restrict the search by precursor mass but instead, compare the query spectrum with *all* spectra in the SVM training dataset. We then assign each compound class individually by a majority vote of the top 5 candidates^17^. Lastly, we evaluated direct class prediction from MS/MS spectra. Here, we employed the kernel support vector machine setup of CSI:FingerID as, to the best of our knowledge, this is currently the best-performing classifier for this task. In short, the molecular fingerprint of CSI:FingerID was replaced by a “ClassyFire fingerprint”, a binary vector where each position indicates the membership of a molecule to a certain compound class. We again ensured structure-disjoint evaluation. We cannot evaluate against propagating class annotations through clustering or molecular networking^27-29^, because such propagation is semi-automated at best and still requires a lot of manual interpretation; see also Fig. 4 below.

**Figure 3.**
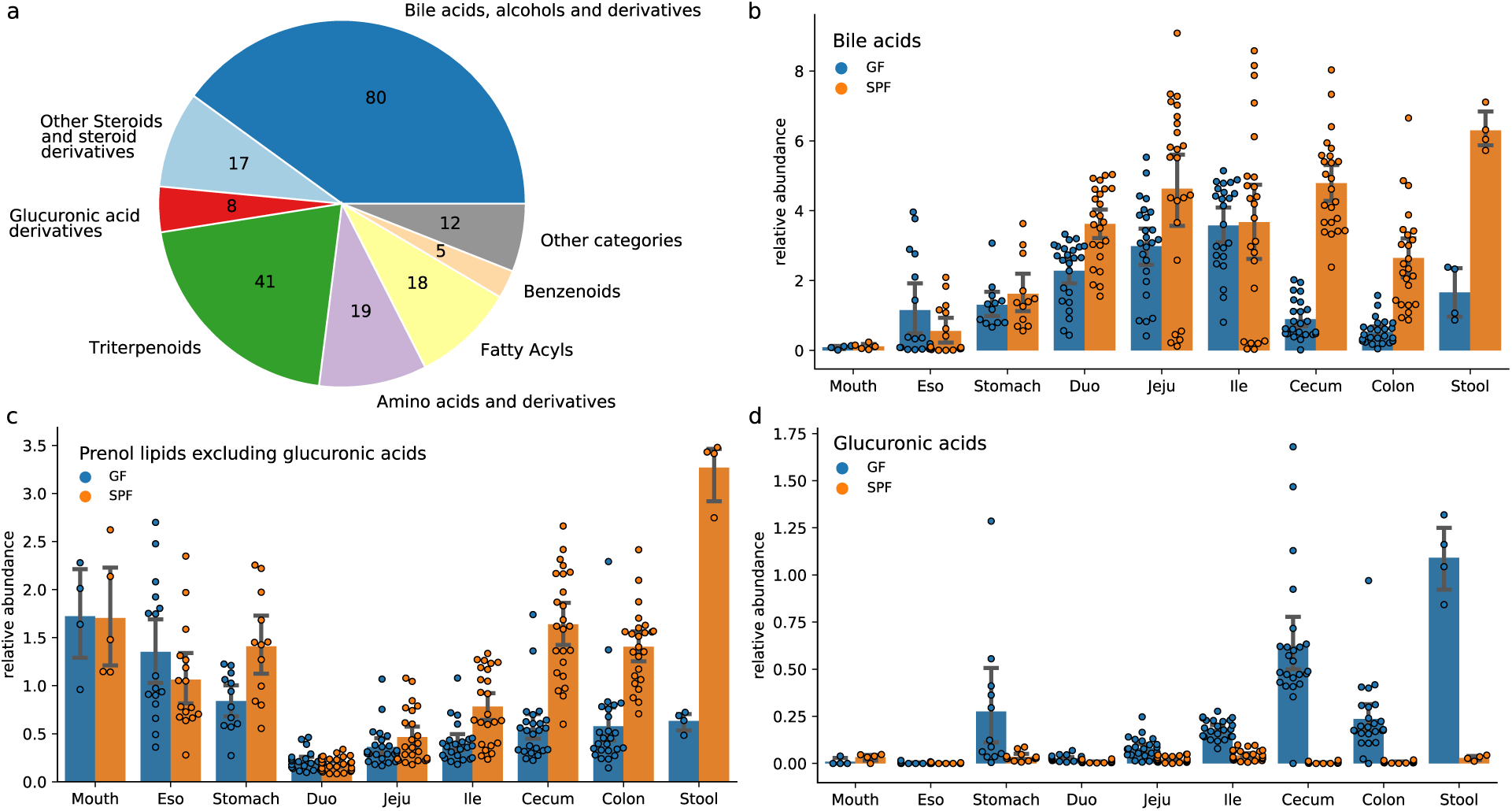
Comparing the digestive system of germ-free and specific-pathogen-free mice. (a) Compound classes of the top 200 most discriminating compounds in the digestive system of germ-free mice (GF) and specific-pathogen-free mice (SPF). (b-d) Summed intensity of all compounds belonging to the class *bile acids* (b), *prenol lipids* excluding *glucuronic acids* (c), and *glucuronic acids* (d) in the digestive system of GF (blue) and SPF (orange). Standard deviation shown as error bars.

**Figure 4.**
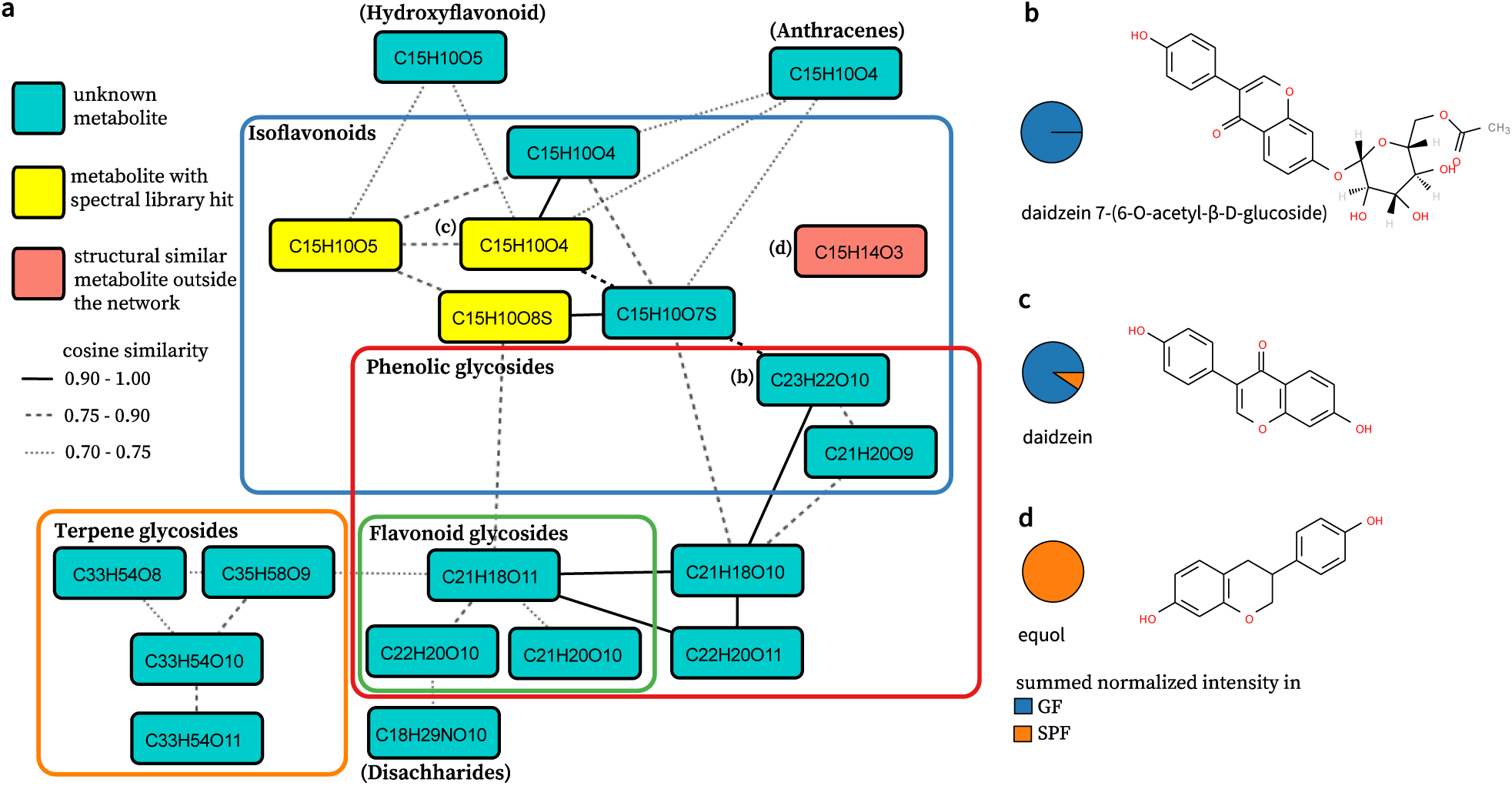
Molecular network of the compound daidzein. Spectral library hits in the network (a) colored in yellow; spectra without spectral library annotation colored in turquoise. Nodes are labeled with the SIRIUS molecular formula annotation. Solid lines are edges in the molecular network with high cosine similarity (0.9 to 1.0), dashed and dotted lines are edges with low cosine similarity (0.75 to 0.9 and 0.7 to 0.75, respectively). CANOPUS compound classifications are indicated as boundaries in the network or written in brackets next to the node. The compound equol (red) is not part of the network but is classified correctly as isoflavonoid by CANOPUS. (b-d) Three compounds were annotated by CSI:FingerID as (b) daidzein 7-(6-O-acetyl-β-D-glucoside), (c) daidzein, and (d) equol. Pie charts show the relative intensity of the compounds in GF samples (blue) and SPF samples (orange) for the large intestine and stool.

CANOPUS clearly outperformed the other methods, as we demonstrate for the independent dataset (Fig. 2b, Supplementary Material 2): The average MCC for rich classes was 0.683 for CSI:FingerID KNN-5, 0.532 for spectral library KNN-5, 0.465 for MetFrag KNN-5, and 0.677 for direct prediction from the MS/MS spectrum (0.729 for CANOPUS). The microaveraged MCC for sparse classes was 0.505 for CSI:FingerID KNN-5, 0.338 for spectral library KNN-5, 0.315 for MetFrag KNN-5, and 0.493 for direct prediction (0.531 for CANOPUS).

### Predicting compound classes without MS/MS training data

CANOPUS can predict compound classes for which no training MS/MS spectra exist at all. To demonstrate this, we re-trained the SVM battery such that all 491 MS/MS spectra of *flavonoid glycosides* were removed (Supplementary Figure 2, Supplementary Material 1). CANOPUS was still able to predict this compound class with MCC 0.861, compared to 0.922 when *flavonoid glycosides* were contained in the MS/MS training data. Similarly, the MCC for the parent class *flavonoid* was 0.850 when *flavonoid glycosides* are absent from training data, compared to 0.880 otherwise. In both cases, the drop of performance is surprisingly small, and the classifiers without training MS/MS spectra still reached outstanding performance.

By concept, direct prediction is not able to predict compound classes without any MS/MS training spectra. Beyond that, the lack of training spectra for one particular subclass can also substantially affect the performance of the parent class predictor. To demonstrate this, we trained a kernel SVM for predicting *flavonoids* from MS/MS data but, again, left out all 491 spectra of *flavonoid glycosides*. We then evaluated the *flavonoid* predictor on the *flavonoid glycosides* MS/MS spectra. Only 6 % of the compounds were correctly classified as *flavonoids*. For comparison, CANOPUS correctly recognized 85% of the *flavonoid glycosides* as *flavonoids*, although the CANOPUS MS/MS training data did not contain a single *flavonoid glycoside* spectrum (Supplementary Figure 2, Supplementary Material 1).

We repeated the above analysis on a second compound class: *Bile acids, alcohols and derivatives*. We trained CANOPUS and an SVM classifier without the MS/MS data of 127 *bile acids*. We found that CANOPUS still shows good prediction performance for *bile acids* (MCC 0.534, compared to MCC 0.757 when *bile acids* are part of the MS/MS training data). The MCC of the superclass remains almost constant (0.852 and 0.898, respectively). CANOPUS correctly predicts 90% of the left-out *bile acid* spectra as *steroids*; in comparison, direct prediction recognized only 57 % of those spectra as *steroids*.

### CANOPUS and metabolomics data analysis

Metabolomics aims at establishing changes in the metabolite profile between different experimental conditions, time points etc. Today, these changes are usually monitored at a “per feature” level; but doing so cannot uncover complex changes in metabolite profiles, just like a complex trait usually cannot be attributed to a single genetic variant. We now demonstrate how monitoring differences on a compound class level allows for a more comprehensive view of the biological system. For that, we re-analyzed the data from Quinn et al.^30^, where tissue samples from different organs of germ-free (GF) and specific-pathogen-free (SPF) mice were measured by non-targeted LC-MS/MS. This study led to the discovery of novel conjugated bile acids in the digestive system of SPF absent in GF mice, and found that these conjugated bile acids are produced by gut microbes that reside in the duodenum, jejunum and Ilium region in healthy mice.

We sorted metabolites by fold change in intensity between GF and SPF samples. MS/MS spectra of each metabolite were classified using CANOPUS; then, classes were statistically tested for overrepresentation (Mann-Whitney *U* test). The most significant compound classes are *Bile acids, alcohols and derivatives* (p-value 3.39 × 10^−104^) as well as its subclasses and parent classes. Other highly significant classes are *triterpenoids* (5.86 × 10^−74^) and *glucuronic acid derivatives* (p-value 5.36 × 10^−27^) (Supplementary Table 1). See Fig. 3a for the classes of the 200 most discriminating compounds.

We found that the abundance of *bile acids* is similar for GF and SPF in the small intestine, but substantially different in the large intestine (cecum, colon) and in stool (Fig. 3b), consistent with the findings of Quinn et al.^30^. Our results also showed that *glucuronic acids* (includes related saccharides) were among the most discriminating compounds; most of them were also classified as *prenol lipids* and *isoflavonoids*. In SPF, non-glycosylated *prenol lipids* appeared to be increasing through the digestive system from stomach to stool (Fig. 3c). In contrast, the abundances of *prenol lipids* in GF did not change notably through the digestive system. For the *glucuronic acid derivatives* class, we observed an opposite trend: these had a relatively lower abundance in SPF and did not show a noticeable trend, but accumulated through the digestive system of GF with highest abundance in stool (Fig. 3d). These results suggest the involvement of the microbiota in the metabolization of the glycosylated *prenol lipids* by cleaving off the sugar acids. To check this hypothesis, we considered two glycosylated compounds detected in GF but undetectable in SPF: We searched for the deglycosylated derivatives in both GF and SPF. Using CSI:FingerID, we annotated the first of these compounds as a glycosylated derivative of the *isoflavone* genistein. The glycosylated derivative was detected only in the GF samples, whereas genistein was detected in both SPF and GF samples. The second compound was annotated as glycosylated daidzein. We found that daidzein and its glycosylated derivative daidzein 7-(6-O-acetyl-β-D-glucoside) are both abundant in GF but undetectable or low abundant in SPF (Fig. 4b-d), which appears to contradict the above hypothesis. However, daidzein is known to be metabolized into equol^31^ and, indeed, equol was detected only in the SPF samples (Fig. 4d). Unfortunately, such an in-depth verification is not possible in general, as for many compounds, the molecular structure cannot be confidently annotated.

Molecular networks are a popular method for pushing the boundaries of database search by propagating annotations via spectral network similarity^32^. The underlying idea is that spectral similarity often implies structural similarity, so that annotations from spectral library search can be propagated through connected subnetworks^28^. For the mouse samples we find 344 molecular subnetworks with at least three compounds, but only 92 of these subnetworks have at least one spectral library match enabling subnetwork annotation; an additional 376 compounds are *singletons* and do not cluster with any other compound in the samples. Furthermore, propagation of compound classes to the full subnetwork, as proposed in ref. ^16^, can lead to partial or imprecise annotations. As an example, see the molecular subnetwork containing the compound daidzein (Fig. 4a). Spectral library search allowed us to annotate structures for four nodes of the molecular network, all of them *isoflavonoids*. But inferring that all other compounds in this subnetwork are *isoflavonoids*, too, is most likely incorrect: CANOPUS annotated *flavonoids* and *terpene glycosides* in this subnetwork. Furthermore, most of the compounds in the network were annotated as *glycosylated compounds* (either *phenolic glycosides* or *terpene glycosides*) by *CANOPUS*, whereas none of the spectral annotations belong to these classes. While daidzein 7-(6-O-acetyl-β-D-glucoside) and daidzein are part of the same network, equol, their metabolic product, is a singleton and is not contained in the network as the MS/MS spectrum itself is quite different and therefore does not align (Fig. 4a). CANOPUS correctly annotated two additional *isoflavonoids* that form singleton subnetworks and, hence, are missing from the daidzein network; these *isoflavonoids* were annotated by CSI:FingerID as acacetin and formononetin, which are structurally similar to daidzein. On a larger scale, we find that CANOPUS compound class annotations and molecular networks usually agree well, see Supplementary Figures 3 and 4.

### Comparative metabolomics of *Euphorbia* species

We use CANOPUS to study the chemical diversity among representative plant species of *Euphorbia* subgenera^20^. This time, we are not considering the fold change of compounds or compound classes, but rather the change of metabolite diversity: How many compounds of a certain class do we find in each sample? To answer this question, it is crucial to annotate the entire set of molecules detected, as done by CANOPUS. Annotating only a small subset by, say, spectral library search will often result in misleading findings: Neither is the identified subset an unbiased subsample of the complete metabolome, nor is the spectral library unbiased for compound classes. Ernst et al.^20^ classified around 30% of the detected compounds, using a combination of spectral library search with GNPS^2^, in-silico search with network annotation propagation^28^, MS2LDA^33^, CSI:FingerID, and also manual inspection. The study showed that the diversity of Euphorbia *diterpenoids*, a type of bioactive compounds actively studied for their antiviral and drug-resistance reversal properties^34^, was larger in the *Esula* and *Euphorbia* subgenus than in the *Chamaesyce* and *Athymalus* subgenus. This change in diversity agrees with the geographic co-location of these plants with certain herbivores, given that *Euphorbia* diterpenoids are known feeding deterrent.

Re-analyzing this dataset with CANOPUS allowed us to reproduce the above biological findings, but also to derive new findings. Different from Ernst et al.^20^, compounds were not aligned across *Euphorbia* species but each species was annotated individually. For each species, we counted the number of compounds belonging to each class, but ignored peak intensities of compounds. First, we consider the seven compound classes investigated in ref. ^20^, see Fig. 5a. For each class, our workflow detected and annotated substantially more compounds than the original study. For *diterpenoids* and *triterpenoids*, we observe a similar class distribution pattern: The *Esula* and *Euphorbia* subgenera have a larger diversity of diterpenoids than the other two subgenera (Fig. 5a, Supplementary Figure); the species from the two subgenera *Euphorbias* and *Athymalus* that are mainly distributed in dry regions, have a slightly higher variety of triterpenoids (Supplementary Fig. 7). *Cholestane steroids* and *steroid lactones* do not show a notable distribution pattern here or in ref. ^20^.

**Figure 5.**
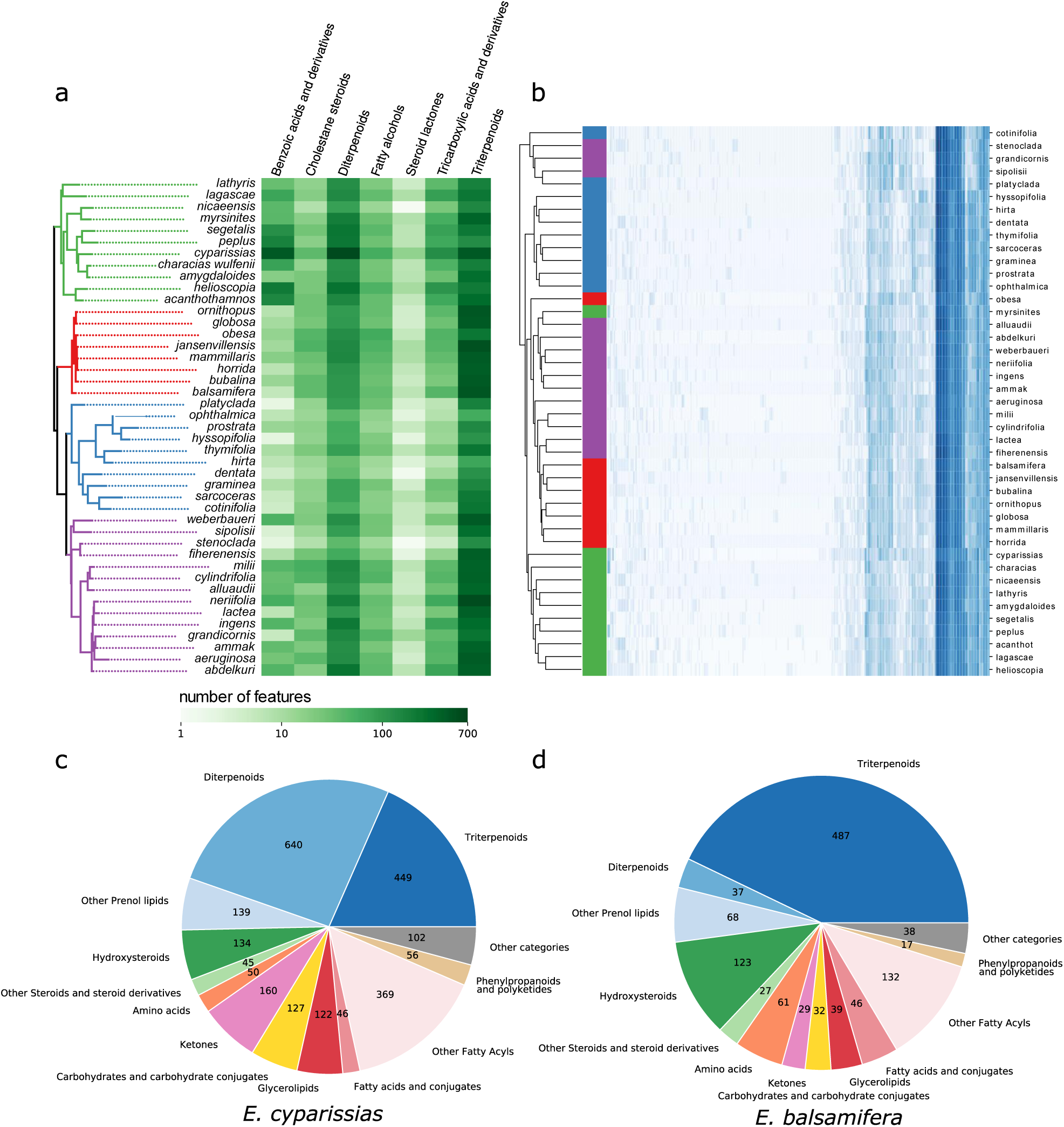
Compound class distribution in euphorbia species. (a) Heatmap of the number of compounds for seven compound classes in the euphorbia species, grouped in a phylogenetic tree computed from genomic data. (b) Clustered heatmap with dendrogram showing hierarchical clustering on compound class distributions. Color on the left indicates the clade a row belongs to: *Chamaesyce* (blue), *Euphorbia* (violet), *Athymalus* (red) and *Esula* (green). (c-d) Distribution of compound classes in *E. cyparissias* (c) and *E. balsamifera* (d). Numbers within the pie charts are the absolute number of compounds of the class annotated in this species.

For *benzoic acid esters*, we observe a notable distribution pattern: These show large variety for the *Esula* and medium variety for the *Euphorbia* subgenus, and no notable variety for the other two. This is in sharp contrast with the findings by Ernst et al.^20^ where one to three *benzoic acids* were found across all species, corresponding to no notable variety. Many compounds annotated as *diterpenoids* were also annotated as *benzoic acid ester, fatty acid esters* or *dicarboxylic acids* (Supplementary Fig. 8). This is characteristic for *Euphorbia* “lower” diterpenes that are often esterified with various acyls^34^. The *Esula* subgenus shows the highest variety of *benzoic acid esters*, followed by the *Euphorbia* subgenus. For the *E. cyparissias* species, 300 of 500 compounds annotated as *diterpenoids* were also annotated as *benzoic acid esters*. This agrees well with observations by Yang et al.^35^ on *E. esula*, a species very closely related to *E. cyparissias*^*36*^. Diterpenoids in *Euphorbia* species are frequently observed with benzoyloxy substituents^34^; yet, it was not known that the occurrence of this acylation differs to this extend for different *Euphorbia* subgenera.

Next, we used hierarchical clustering to cluster *Euphorbia* species based on CANOPUS class annotations (Fig. 5b). The resulting chemodendogram shows good agreement with a phylogenetic tree of *Euphorbia* computed from genomic data^20, 37, 38^: The subgenera *Esula, Athymalus* and *Chamaesyce* form almost perfect clades in the chemodendogram. In contrast, the clustering from ref. ^20^, computed from a spectral similarity-based distance metric^39^, shows little similarity to the phylogeny.

Exploring the distribution of compound classes allowed us to spot differences and common features between samples (Supplementary Material 3). Consider *E. cyparissias* (Fig. 5c) and *E. balsamifera* (Fig. 5d): The variety of *diterpenoids* observed in *E. cyparissias* (640 *diterpenoids*) was higher than in *E. balsamifera* (37 *diterpenoids*). CANOPUS annotated 9 and 17 compounds as *bile acids* in *E. cyparissias* and *E. balsamifera*, respectively. It is noteworthy that these are no annotation errors: The ClassyFire ChemOnt ontology does not distinguish between *bile acids* and *phytosteroids*.

## Discussion

CANOPUS is an automated method for the systematic classification of compounds from fragmentation spectra. CANOPUS predicts 1,270 compound classes and is best-of-class for this task. Surprisingly, CANOPUS can reliably predict numerous compound classes, even for which little or no MS/MS training data are available, and is not distracted if MS/MS data is available only for a subclass. Its integration into SIRIUS allows us to perform the entire workflow from feature detection to compound classification in one tool and on complete datasets. CANOPUS can also import data from popular mass spectrometry frameworks such as MZmine^40^, OpenMS^41^, and XCMS^42^.

We demonstrated how to use CANOPUS for comparative metabolomics applied to microbiome research and chemotaxonomical investigations in plant. Class annotations allowed us to infer novel biological findings, without the need for annotating all MS/MS spectra with a spectral library or structure database. Findings include that gut microbiota are likely involved in the metabolization of glycosylated lipids such as plant saponins, and characteristic distribution pattern for benzoic acid esters in diterpenoids across the *Euphorbia* subgenera. CANOPUS classifications can be used as part of semiquantitative and qualitative metabolomics data analysis; in particular, class distributions allow us to compare samples or species that have little or no overlap in compounds. We anticipate that in the foreseeable future, a large fraction of compounds detected by non-targeted mass spectrometry will remain without structural annotation; CANOPUS can classify these unannotated fragmentation spectra and allows us to directly deduce information from compound class distribution. In the future, CANOPUS can predict classes outside the ClassyFire ontology, and other chemical properties for which few or no MS/MS training data are available; examples are the prediction of antiobiotic activity or toxicity of compounds.

## Methods

### Training and evaluation data

For training the kernel support vector machines (SVMs) we use mass spectral data from 23,965 compounds with 16,701 unique 2D structures; 13,912 compounds are from NIST 2017 (commercial; NIST, v17), 7,680 compounds are from GNPS^2^, and 2,373 compounds are from MassBank^1^. This MS/MS dataset is referred to as *SVM training dataset*, to differentiate it from the structure dataset used to train CANOPUS. As an independent dataset, we use mass spectral data from 3,385 compounds in the “MassHunter Forensics/Toxicology PCDL” library (Agilent Technologies, Inc., Santa Clara, USA). We use only compounds measured in positive ion mode, as these are more abundant.

The training set for the deep neural network (DNN) consists of molecular structures from numerous public databases like KNApSAcK^43^, HMDB^44^, KEGG^45^, UNDP, and others. We excluded all structures from the DNN training dataset that were contained in the SVM training dataset or the independent dataset. In total, the DNN training set consists of 1,106,938 structures along with their ClassyFire compound classes and molecular fingerprints.

### Chemical Classes

The ChemOnt ontology of ClassyFire consists of 4,825 classes^13^ which are organized in a tree. In practice, every compound is assigned to several classes. For many classes, we do not find a single positive example in any biological structure database. We use 1,270 classes for which at least 500 example structures are present in the DNN training set. In the future, we will classify larger parts of PubChem to increase the number of training data and, thus, also the number of trainable classes.

Historically, biomolecules have been classified based on common biosynthetical origin, or chemical characteristics. However, doing so, automated classification of compounds has been a non-trivial problem even if the compound structure is known^46^. In contrast, ClassyFire classes do not depend on biological precursors or characteristics but instead, can be deterministically computed from the molecular structure. This allows us to assign each class for every compound in a structure database.

### Molecular fingerprints and fingerprint prediction

As described by^10, 26^, we use molecular properties from several known molecular fingerprints: Namely, CDK Substructure fingerprints, PubChem CACTVS fingerprints, Klekota-Roth fingerprints^47^, FP3 fingerprints, MACCS fingerprints, and Extended Connectivity Fingerprints^48^. Molecular properties are computed from molecular structure using the Chemistry Development Kit (CDK) 2.1.1^49^. In addition, we use 490 molecular properties that describe larger substructures and will be added to SIRIUS and CSI:FingerID in an upcoming release. For CANOPUS, the additional molecular properties did not result in an improved prediction performance, so we leave out the details. We discard molecular properties with less than 20 positive examples and that cannot be predicted with reasonable quality (F1 above 0.25) during cross-validation. In total, CSI:FingerID predicts 3,609 molecular properties.

We find that building molecular structures from InChI (IUPAC International Chemical Identifiers) results in inconsistent representations of molecules, as structures are not standardized. Hence, we now build molecular structures from PubChem canonical SMILES (Simplified Molecular Input Line Entry Specifications) in all cases^50^.

Training the CSI:FingerID kernel SVMs is done as described in^26^. We train ten models in tenfold cross-validation, such that every model is trained on 90% of the structures. Cross-validation is performed structure-disjoint: Whenever we predict the molecular fingerprint of a query compound in evaluation, we use the cross-validation model that has not seen the query structure during training. We do so for both the SVM training dataset and the independent dataset. To this end, all compounds are novel, in the sense that none can be identified by dereplication (spectral library searching).

### Predicting compound classes from molecular fingerprints

Assume that we know the exact molecular fingerprint and molecular formula of a query compound, but not its structure; can we predict whether it belongs to a certain compound class? We use a Deep Neural Network (DNN)^22^ for this purpose, simultaneously predicting all compound classes. DNNs can be trained with millions of training examples in reasonable time. Molecular formulas are encoded as feature vectors containing the number of atoms for each element, the mass, the RDBE value, as well as the ratios between some elements (see Supplementary Table 2 for all features). The binary molecular fingerprint and the molecular formula features constitute the input of the DNN.

The above DNN appears to be of no practical use: To compute the exact fingerprint of a compound, we have to know its molecular structure; but if we know the molecular structure, there is no need to apply the DNN, because we can deterministically find the correct answer using ClassyFire. The trick is that we can use a *predicted fingerprint* as input of the DNN: We use the probabilistic molecular fingerprint predicted by CSI:FingerID as well as the molecular formula computed by SIRIUS (see below) as input. The predicted fingerprint can either be transformed to a binary fingerprint, or we can directly use the probabilistic fingerprint as input of the DNN. We found that the latter option performs substantially better (data not shown).

### Simulating probabilistic fingerprints

It turns out that we can improve the performance of the above DNN as follows: During training of the DNN, we use binary fingerprints (computed from structure) whereas in application, we use a probabilistic fingerprint predicted from the query MS/MS spectrum; the later fingerprint contains errors and uncertainties. To improve the DNN performance, we present two probabilistic methods to introduce errors and uncertainties into the training fingerprints. Due to the probabilistic nature of the methods, one molecular structure will result in different probabilistic fingerprints, all of which we can use to train the DNN.

The first method of sampling probabilistic fingerprints considers all molecular properties individually: For each molecular property *i*, we have trained an individual SVM to predict the property from MS/MS data. We record all Platt probabilities that were estimated for positive property *i*: Let 𝒫_*i*_ be the set of all Platt probability estimates in cross validation, for all MS/MS spectra in the training set where the SVM should have predicted a positive outcome for property *i*. Analogously, we record Platt probability estimates for negative property *i* in 𝒩_*i*_. Given a compound from the structure database, we know its exact binary fingerprint. Consider a molecular property *i*: If the property is present in the compound, we can uniformly sample from 𝒫_*i*_ to simulate a Platt probability; if is not present, we uniformly sample from 𝒩_*i*_. But this may result in overfitting of the DNN, as we are using exactly the same real numbers for training which are later used for evaluation. To this end, we also sample from the “holes” between values in 𝒫_*i*_: We sort the set 𝒫_*i*_ := {*p*_1_, …, *p*_*n*_} such that *p*_1_ ≤ … ≤ *p*_*n*_, and assume *p*_0_ := 0 and *p*_*n*+1_ := 1. We uniformly draw a random number *x* ∈ (0, *n* + 1) and then interpolate between values *p*_*k*_ and *p*_*k*+1_, with *k* := ⌊*x*⌋. That is,

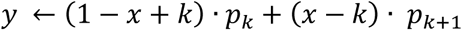

is the simulated Platt probability. Analogously, we can do so for 𝒩_*i*_. This can be interpreted as drawing random numbers using a kernel density estimate of the observed Platt probabilities; we avoid elaborate kernel estimates to guarantee swift running times.

The disadvantage of drawing every position independently is that we completely ignore correlations between fingerprint positions. If two molecular properties are highly correlated with each other, we can assume that prediction errors correlate, too. To this end, our second sampling method draws multiple positions at once. First, we define further sets of Platt probabilities for each position in the fingerprint, namely *TP*_*i*_, *FP*_*i*_, *TN*_*i*_, and *FN*_*i*_. These are defined analogously to 𝒫_*i*_ and 𝒩_*i*_, but contain Platt estimates for true positives, false positives, true negatives and false negatives for molecular property *i*. Let ℱ be a binary fingerprint from the structure database for which we want to simulate a probabilistic fingerprint. We measure the similarity of two fingerprints using the Tanimoto coefficient (Jaccard index)^51^. We first sort all structures from the SVM training dataset in descending order of their Tanimoto coefficient to ℱ. We then pick the *k*-th structure and its binary fingerprint *B*, together with its predicted probabilistic fingerprint 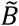. Here, *k* is a random number drawn from a geometric distribution with small parameter *p*; here, we use *p* = 0.2. For each molecular property *i* with *B*_*i*_ = ℱ_*i*_, we randomly draw a Platt probability from the appropriate set *TP*_*i*_, *FP*_*i*_, *TN*_*i*_, and *FN*_*i*_, using the procedure described above. For example, if 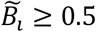 and *B*_*i*_ = 1, then this is a true positive prediction and we sample from *TP*_*i*_; if 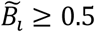 and *B*_*i*_ = 0, then this is a false positive prediction and we sample from *FP*_*i*_, and so on. The remaining positions differ between ℱ and *B*; for these, we repeat the above procedure: This time, we only use the subset of remaining positions to calculate the Tanimoto between ℱ and the structures from the SVM training data and for sampling the probabilities.

We can think of the second sampling method as sticking together the simulated fingerprint using parts of probabilistic fingerprints available for training. One may assume that this sampling method yields “more realistic” probabilistic fingerprints. On the other hand, it may lead to probabilistic fingerprints which are “too similar” to the training data and, thus, may result in overfitting of the DNN. As both sampling strategies have advantages and disadvantages, we do not decide for one, but simulate fingerprints using the first and the second sampling strategy alternately.

### DNN architecture and training

Both input layers are centered feature-wise: For centering the molecular fingerprint, we calculate the mean for every predicted molecular property in the SVM training dataset. We stress that no structures from the SVM training dataset are used for training the DNN, but only statistics about these structures such as the mean of each molecular property, as this is needed for centering the feature vectors; we also use the SVM training dataset for determining the early stopping of the DNN training. Recall that no structures from independent dataset are used for training the DNN, either. The molecular formula feature vectors are normalized such that every feature has unit variance; we use all molecular formulas from the DNN structure training database to calculate the empirical mean and variance.

Instead of concatenating both input layers directly, we first connect the fingerprint input layer with a fully connected inner layer of 3,000 neurons, and the molecular formula input layer with a fully connected inner layer of 16 neurons (see Fig. 6). The outputs of both inner layers are concatenated and fed into two additional inner layers with 3,000 and 5,000 neurons. All inner layers are using the ReLu activation function, a learnable bias, an *L*_2_ regularization as well as a dropout of 50 % of the neurons^52^. The output layer is linear and predicts 1,270 compound categories. We use the sigmoid cross-entropy as loss function and Adam as optimizer^53^. The DNN training is performed using the TensorFlow library^54^. Training is done in minibatches of roughly 6,000 structures; we draw compounds from the pool of all available training compounds such that every compound class is present in each minibatch at least one time. The sign of each output neuron encodes whether a compound belongs to the respective compound class. We stop training as soon as the MCC on the SVM training dataset did not improve anymore; this usually happened after about 15,000 iterations. We do not report epochs, as our training data is randomized.

**Figure 6.**
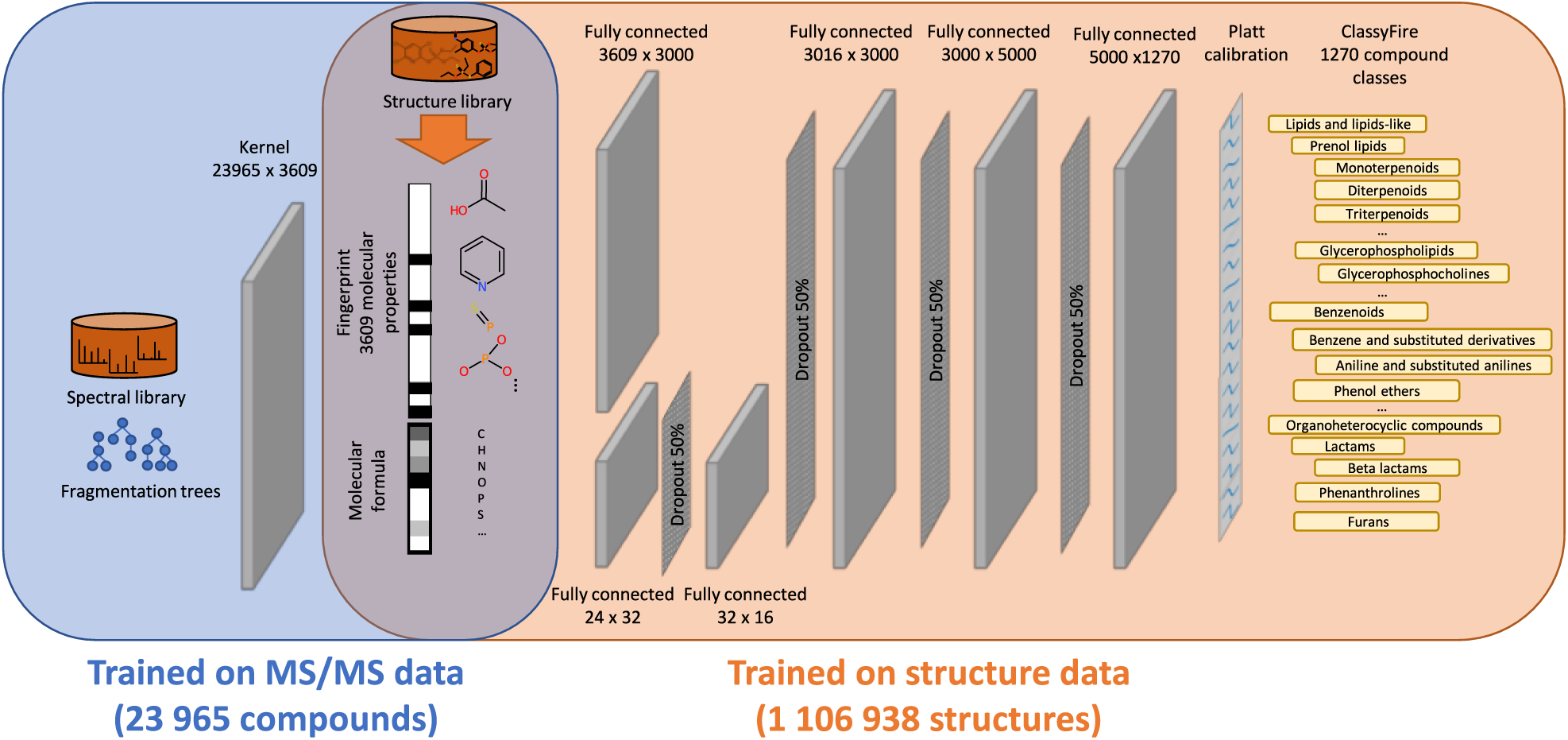
Heterogeneous training for compound class prediction. First, a battery of kernel SVMs is trained on spectral training data (blue area left) to predict the molecular fingerprint. Second, a DNN predicts compound classes from the molecular fingerprint (orange area right) and the molecular formula. The network consists of several fully connected layers with ReLu activation function, and dropout layers in-between. Platt calibration is used to transform the network output to posterior probabilities. The main trick is that the molecular fingerprints used for training the DNN are computed directly from the structure database, without the need of spectral training data.

The following is not performed as part of our method evaluation, but only when applying the method to biological data: We transform the linear outputs of the DNN into posterior probabilities using Platt calibration^55^. For learning the parameters of the logistic function, we use the predicted fingerprints from the SVM training dataset. If we do not have enough positive examples (less than 30 examples), we add simulated probabilistic fingerprints. After the calibration, we update the weights of the network a last time, using the SVM training dataset as input.

### Assigning molecular formulas

Using the molecular formula as input of the DNN requires us to first identify the correct molecular formula. But this is necessary anyways, as the kernel support vector machines from ref. ^10^ operate on fragmentation trees, which are an outcome of the molecular formula identification with SIRIUS^56^.

As our method is targeting novel compounds, we must not assume that the molecular formula of the query compound is recorded in any molecular structure database. We initially use both isotope patterns and fragmentation patterns to determine the molecular formula using SIRIUS 4^26^. Unfortunately, molecular formula identification rates severely drop for compounds above 500 Da, see Fig. 5 in ref. ^56^. ZODIAC improves molecular formula annotations of complete LC-MS runs using a network-based approach where compatible molecular formula assignments support each other, and assignments for the complete dataset are estimated by Gibbs sampling; in evaluations, ZODIAC reaches substantially smaller error rates than SIRIUS^57^. We add all reference compounds from GNPS, MassBank and NIST as anchors into the ZODIAC network.

### Method evaluation

We evaluate against four other methods for compound class assignment: *direct prediction, k-Nearest Neighbor* (k-NN) using either MetFrag^24^ or CSI:FingerID^10^ as the underlying search engine, and k-NN searching in a spectral library.

#### Direct prediction

We can think of the compound classes as additional molecular properties of a compound, and directly predict the corresponding fingerprint from MS/MS data. This approach has been suggested repeatedly in the literature, in particular for Gas Chromatography (GC) MS data, but usually targeting compound classes defined by the presence or absence of a certain substructure. Here, we employ the kernel SVM machinery behind CSI:FingerID for direct prediction, following the usual CSI:FingerID training and evaluation setup. We argue that this setup is currently best-in-class for direct prediction. By design, direct prediction cannot predict compound classes for which there is no positive MS/MS training data; it cannot predict compound classes if training data are available for a subclass only, as the resulting predictor is in fact targeting the subclass; and, as we do not distinguish between compounds with known structure and compounds with MS/MS data, it is not possible to evaluate if direct prediction will show reasonable performance for truly novel compounds with unknown structure.

#### k-Nearest Neighbor (k-NN)

MetFrag and CSI:FingerID are methods for compound structure identification by searching MS/MS spectra in some structure database such as PubChem. Given an MS/MS spectrum, both tools report a ranked list of structure candidates. This approach must fail for novel structures that are not contained in structure databases. However, for the task of compound classification it is sufficient that the top-ranking candidates belong to the same compound class as the query compound. The compound classes for any given candidate structure are computed using the ClassyFire webservice (http://classyfire.wishartlab.com/). A k-NN uses the compound class annotations of the top *k* candidates and, for each compound class, decides via majority vote (yes/no) of the top *k* candidates whether the class is assigned to the query. We search in the PubChem structure database, considering all structures with the same molecular formula as the query, and use MetFrag or CSI:FingerID, as the search engine. We removed structures in the independent dataset from PubChem and the SVM training dataset before searching, as it was done for training the DNN. This corresponds to the situation that the measured compound is a novel structure. We evaluated parameters k=1 and k=5, and found that k=5 performs slightly better for both MetFrag and CSI:FingerID (data not shown).

It is noteworthy that this approach has a number of conceptual shortcomings. We noted that if there exist no molecular structures with the query molecular structure, then the method cannot assign any compound classes. This shortcoming is the most obvious but only indicates an underlying issue, resulting in numerous similar problems: As an example, for *k*=5, we need at least three positive examples with this molecular formula so that a compound class can be positively assigned.

Presented an outlier structure substantially different from all other structures with the same molecular formula, we will predict all corresponding compound classes wrong *even if the correct structure is part of the structure database we search in*. More generally, if a certain compound class is over- or underrepresented for a particular molecular formula, then it is more likely to be predicted “yes” or “no”, respectively, independent of the actual data. These shortcomings do not result in bad evaluation statistics, though: Considering any *reference compound* used for evaluation, not only its structure is present in PubChem, but also several structures with high similarity and identical molecular formula. To this end, *evaluation statistics of the k-NN classifiers must be interpreted with care*. It is also noteworthy that the abovementioned issues are *not* the ones usually attributed to k-NN classifiers (curse of dimensionality, label noise).

Instead of searching in a structure database, we can also compare the query spectrum against spectra in a spectral library, and obtain a ranked list of reference spectrum candidates. Considering the small size of the spectral library, we do not filter by the precursor mass of the query. For comparing two spectra, we use the cosine similarity of the two spectra, as well as of the inverse spectra (obtained by subtracting each peak mass from the precursor mass) and sum up both values. Using the spectrum and its inverse allows for comparing spectra of compounds that differ in mass but are structural very similar^14, 58^. We search in the SVM training dataset as our spectral library. For each query, we removed all spectra from the spectral library corresponding to the query structure. We found that hybrid search works better than just comparing spectra by their cosine (data not shown). All of the abovementioned issues also apply to the spectral library k-NN, considering the size of the spectral library; in addition, it suffers from all issues of direct prediction.

#### Matthews correlation coefficient

The Matthews correlation coefficient (MCC) is defined as

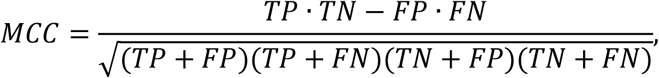

where *TP, TN, FP, FN* are the numbers of true positives, true negatives, false positives and false negatives of a binary classification, respectively^23^. The MCC lies between -1 and +1, where +1 corresponds to a perfect classification, 0 to a random classification, and -1 to a “perfectly wrong” classification. The MCC is considered more informative than F1 score and accuracy, because it takes into account the balance ratios of these four values; in particular, the F1 score of a random classification is non-zero and depends on the actual classification task, rendering predictor performance incomparable. The MCC is equivalent to the Pearson correlation coefficient between observations and predictions.

### LC-MS data processing

For the processing in SIRIUS 4.4, one or more LC-MS/MS runs must be provided in mzML or mzXML format. Feature detection in SIRIUS 4.4 is similar in spirit to a *targeted analysis*: instead of searching for all features in a run, SIRIUS is first collecting all MS/MS spectra and their precursor information. It then searches for features that are associated with those MS/MS spectra: these are the precursor ions, adduct ions and isotope peaks. The precursor information reveals the retention time and mass range where the precursor ions can be detected. Adducts and isotopes can be found using predefined lists of mass differences. Fragmentation spectra assigned to the same feature (precursor ion) are merged. SIRIUS 4 is using the same feature alignment method as MZmine^40^.

### Mice dataset

We analyzed 834 LC-MS runs from MassIVE (id: MSV000079949)^30^. The corresponding samples were taken from different organs of eight mice; four have an intact microbiome (specific-pathogen free, SPF), while four are germ-free (GF). Feature detection and feature alignment were performed using SIRIUS 4.4. We only kept features for which we had MS/MS measurements in at least two samples. We used ZODIAC to improve molecular formula annotation^57^. To speed up running times, only compounds below 860 Da were considered.

This resulted in a feature quantification table, where each row corresponds to compound and each column to its intensity (maximum peak height) per sample. We predicted compound classes for each compound. We subtracted the blank intensities from the table, and performed quantile normalization to make the samples comparable. Next, we analyzed which compound classes have a different abundance between GF and SPF in the digestive system. In the following, we only consider samples from the organs mouth, esophagus, stomach, duodenum, jejunum, ileum, cecum, colon, as well as from stool.

Summing up all intensities over all compounds of a class already gives helpful insights, see Fig. 3b-d. However, this assumes that all compounds of a class change their intensity in the same direction. A better indicator is the fold change, which is calculated as the median of all intensities in GF samples divided by the median of all intensities in SPF samples. A pseudo-count must be added beforehand, to avoid division by zero. We use the 1% percentile of non-zero intensities in the quantification table as a pseudo-count.

We sorted compounds by absolute logarithm (base 10) of their fold changes. Values below 1 (fold changes between 0.1 and 10) were set to zero, as we do not consider such fold changes to be informative. For each compound class, we checked if compounds of this class appear more often in the higher discriminative region; this is done using a one-tailed Mann-Whitney-U permutation test^59^. We then sort compound classes based on their p-value. Compound classes with the lowest p-value belong to classes which trigger the metabolic difference between GF and SPF samples (Supplementary Table 1).

We uploaded quantification table and input MS/MS spectra to GNPS and performed feature-based molecular networking^2, 60^. We downloaded the Cytoscape files for the network visualization. We mapped CANOPUS class annotations to each node in the network. For evaluating results, we compared selected class annotations with GNPS library hits and with CSI:FingerID database search results (searching in PubChem).

### Euphorbia dataset

We use LC-MS runs from the study by Ernst et al.^20^ downloaded from MassIVE (id: MSV000081082). In total, we analyzed samples from 43 different Euphorbia species. Feature detection was performed by SIRIUS 4.4. In contrast to ref. ^20^ we did not align features, because samples were taken from different species and, therefore, have a different metabolic composition; see Fig. 3b in ref. ^20^. For most samples, SIRIUS detected more compounds than ref. ^20^ (Supplementary Fig. 5). For molecular formula annotation we used SIRIUS and ZODIAC^26, 57^. ZODIAC processed the plant samples of all species within one network. Compound classes were predicted for all compounds using the best scoring ZODIAC molecular formula. For counting the number of compounds per class, we considered all compound class assignments with a probability greater than 0.5.

### Chemotaxonomy

We used WPGMA hierarchical clustering for computing the dendrogram in Fig. 5b. As the distance metric, we chose the Euclidean distance over the normalized compound class distribution; this is the vector with the logarithmized number of compounds per compound class plus one, centered to zero mean and unit variance. This turns out to be critical, as the number of detected compounds differ substantially between species (Supplementary Fig. 5). We used the scipy library for clustering^61^.

## Acknowledgment

We gratefully acknowledge financial support by the Deutsche Forschungsgemeinschaft (BO 1910/20) to SB, KD and ML, and of the Academy of Finland (310107/MACOME) to JR. PCD was supported by the Gordon and Betty Moore Foundation (GBMF7622), and the U.S. National Institutes of Health (P41 GM103484, R03 CA211211, R01 GM107550). LFN was supported by the U.S. National Institutes of Health (R01 GM107550), and the European Union’s Horizon 2020 program (MSCA-GF, 704786). We thank F. Kuhlmann, and Agilent Technologies Inc. (Santa Clara, CA, USA) for providing data we used for evaluating CANOPUS. We thank Yannick Djoumbou Feunang, David Arndt, and David Wishart for providing ClassyFire annotations for a database of molecular structures.

## Author contributions

KD, JR, and SB designed the research. KD and SB developed the computational method. KD implemented the computational method with contributions by ML, MF and MAH. MF integrated CANOPUS into SIRIUS 4.4. KD, LFN, and PCD applied and evaluated the method on the biological studies. SB, KD, LFN and PCD wrote the manuscript, in cooperation with all authors.

## Supplementary

**Supplementary Figure 1.**
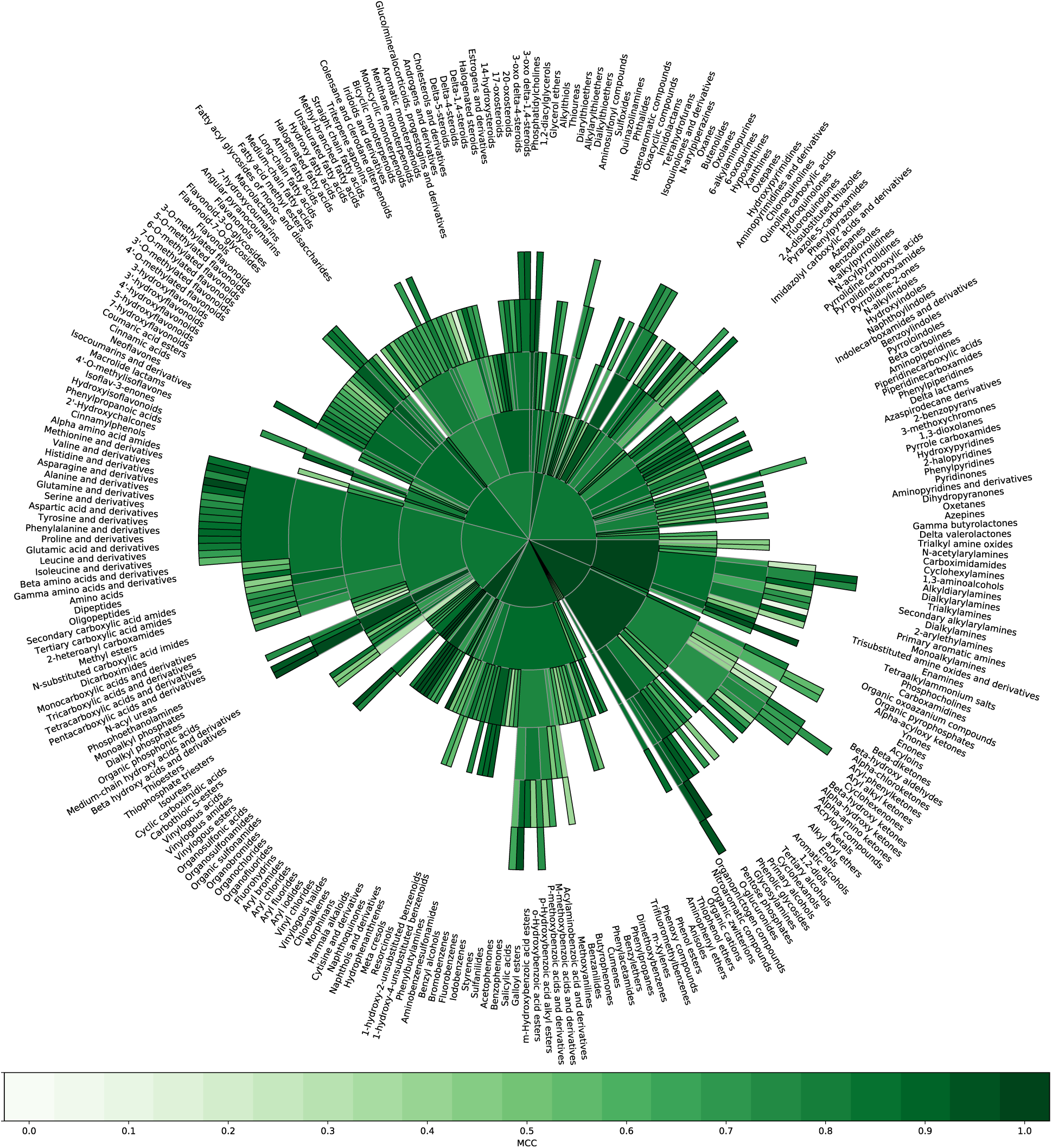
CANOPUS performance sunburst plot. Matthews correlation coefficient (MCC) for the 732 of 1,270 compound classes with at least 50 positive examples. SVM training dataset. A darker green coloring corresponds to better prediction performance for the class. The size of each slice is chosen such that all classes fit into the figure and has no further meaning. Inner slices represent parent classes of outer slices. Two basal classes (*organic oxygen compounds* and *organooxygen compounds*) included in the plot for completeness.

**Supplementary Figure 2.**
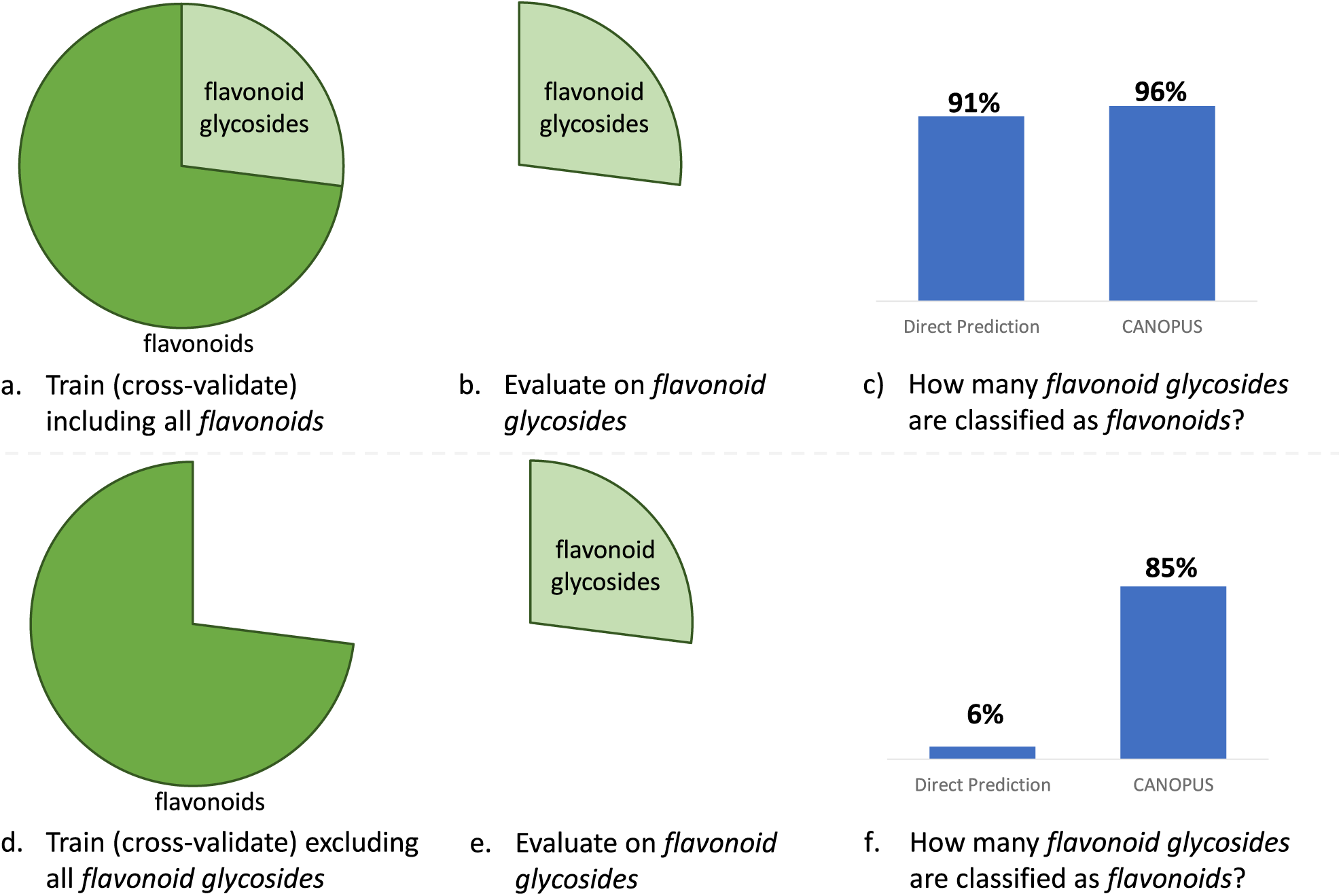
Effect of removing a subclass from the MS/MS training data. (a-c) Regular evaluation setup: classes and subclasses are distributed into cross-validation folds, ensuring that methods are never evaluated on the same MS/MS data or structures they were trained on. (d-f) We remove all *flavonoid glycosides* (the subclass) from the MS/MS training data (d), and then evaluate the predictor for *glycosides* (the class) on these removed MS/MS spectra (e). A perfect method would still classify all *flavonoid glycoside* MS/MS spectra as *glycosides* (f). CANOPUS exhibits only a small drop (85% to 96%) in correct classifications (c,f). In contrast, direct prediction performed mostly on par with CANOPUS before removing *flavonoid glycosides* from the MS/MS training data (c), but misses almost all of them (94%) afterwards (f). We were able to attribute this to the presence of *isoflavonoide glycosides* in the training data; these do not belong to the *flavonoid* class, but have highly similar structures and MS/MS spectra, except for the presence of a sugar residue. We observed that direct prediction in (d-f) uses the presence of a sugar residue to infer that a MS/MS spectrum is not a glycoside. In contrast, CANOPUS does not fall for this “bait”; heterogeneous training allows us to integrate the substantially more comprehensive structure data in its predictions.

**Supplementary Table 1.** The sixty most discriminative compound classes for differentiating between germ-free and specific-pathogen-free mice. Reported p-values are calculated using the one-tailed Mann-Whitney-U test and corrected via Bonferroni-correction. Column ‘#’ contains the number of compounds annotated with this compound class.

| Compound class | p-value | # | Compound class | p-value | # |
| --- | --- | --- | --- | --- | --- |
| Steroids and steroid derivatives | 2.84E-141 | 505 | Oxacyclic compounds | 4.63E-05 | 1046 |
| Cyclic alcohols and derivatives | 3.06E-118 | 616 | Monosaccharides | 4.99E-04 | 289 |
| Bile acids, alcohols and derivatives | 3.39E-104 | 223 | Carboxylic acids | 7.35E-03 | 1179 |
| Hydroxysteroids | 4.57E-98 | 314 | Organosulfonic acids and derivatives | 1.02E-02 | 156 |
| Hydroxy bile acids, alcohols and derivatives | 7.38E-84 | 182 | Organic sulfonic acids and derivatives | 1.02E-02 | 156 |
| Triterpenoids | 5.86E-74 | 113 | Flavonoid glycosides | 2.11E-02 | 12 |
| 3-hydroxysteroids | 2.69E-72 | 173 | Acetals | 2.27E-02 | 451 |
| Prenol lipids | 5.41E-43 | 241 | Sulfonyls | 2.64E-02 | 177 |
| Secondary alcohols | 5.81E-39 | 1651 | 1-benzopyrans | 4.66E-02 | 64 |
| Alcohols and polyols | 1.27E-27 | 2119 | Flavonoids | 7.72E-02 | 17 |
| O-glucuronides | 5.36E-27 | 43 | Isoflavonoids | 8.08E-02 | 10 |
| Sugar acids and derivatives | 5.36E-27 | 43 | Ethers | 9.04E-02 | 1073 |
| Glucuronic acid derivatives | 5.36E-27 | 43 | Thiazoles | 9.55E-02 | 14 |
| Glucuronides | 7.39E-26 | 42 | Cyclohexenones | 1.07E-01 | 17 |
| Beta hydroxy acids and derivatives | 2.06E-23 | 50 | Sulfuric acid esters | 1.43E-01 | 10 |
| Hydroxy acids and derivatives | 2.35E-22 | 49 | Benzopyrans | 2.22E-01 | 70 |
| Steroid esters | 5.43E-15 | 43 | Sulfuric acid monoesters | 3.54E-01 | 11 |
| Ketones | 8.51E-15 | 172 | Oxanes | 3.56E-01 | 505 |
| Pyrans | 2.14E-13 | 90 | 5-hydroxyflavonoids | 5.22E-01 | 16 |
| Terpene glycosides | 8.03E-13 | 27 | Vinylogous acids | 7.89E-01 | 29 |
| Polyols | 8.66E-12 | 502 | Carbohydrates and carbohydrate conjugates | 8.96E-01 | 472 |
| Phenolic glycosides | 1.52E-11 | 16 | Steroidal glycosides | 1.01E+00 | 21 |
| Tetrahydrofurans | 1.46E-10 | 38 | Pyranones and derivatives | 1.33E+00 | 41 |
| Alkanesulfonic acids | 3.20E-09 | 102 | Hydroxyflavonoids | 1.47E+00 | 18 |
| Organosulfonic acids | 3.20E-09 | 102 | Monocarboxylic acids and derivatives | 1.67E+00 | 1487 |
| Chromones | 4.22E-09 | 21 | Flavones | 3.08E+00 | 11 |
| Alkanesulfonic acids and derivatives | 5.15E-09 | 103 | Tertiary alcohols | 3.63E+00 | 92 |
| Oxosteroids | 9.43E-09 | 34 | 7-hydroxyflavonoids | 3.72E+00 | 11 |
| Lipids and lipid-like molecules | 2.49E-05 | 2918 | Hexoses | 5.66E+00 | 69 |

**Supplementary Figure 3.**
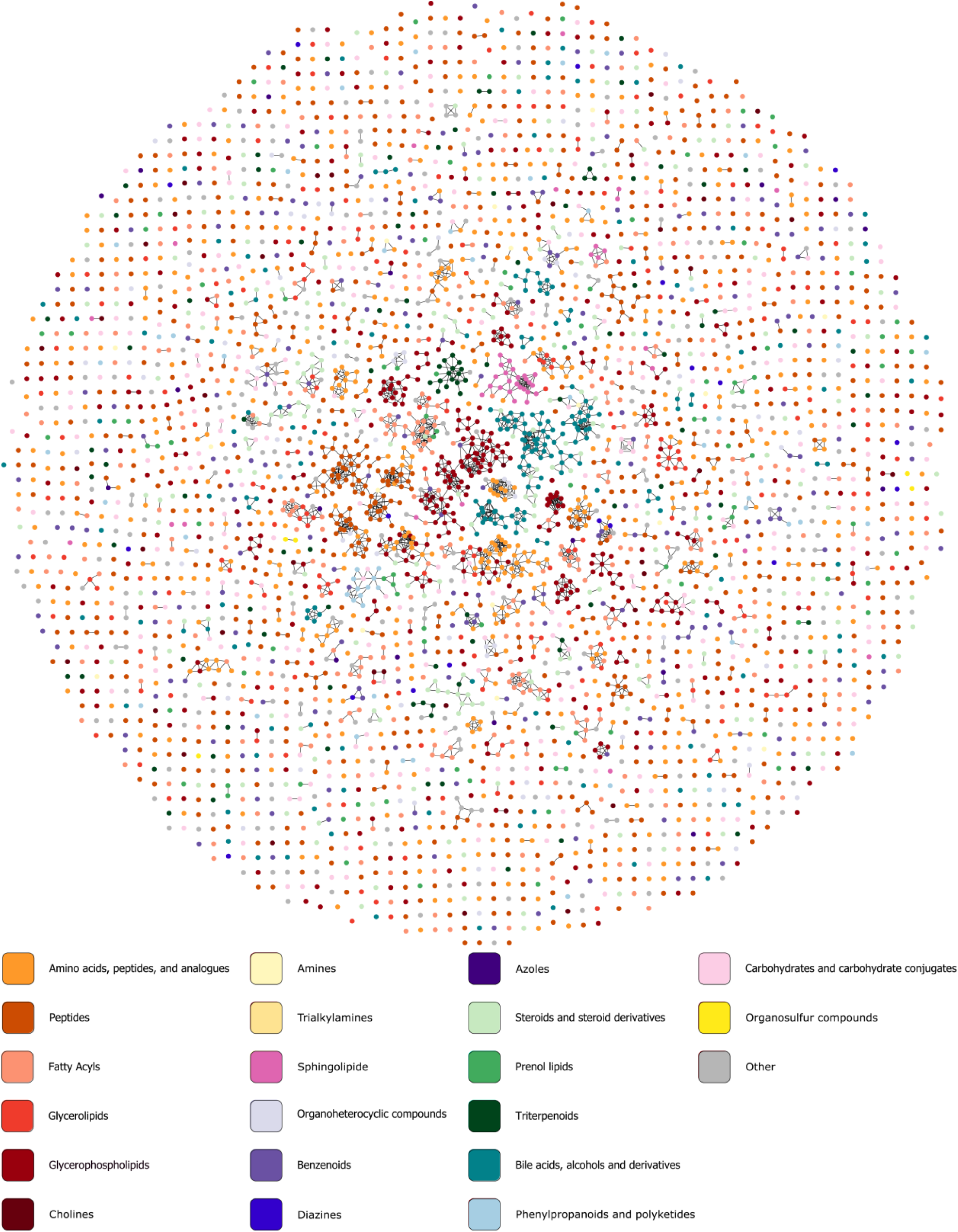
Molecular network and compound class annotations for the mice digestive system. Node colors indicate the compound class annotated by CANOPUS; displayed compound classes were manually selected. When a compound is annotated with multiple classes, the class with the larger structural pattern is selected. Nodes are connected by an edge if the spectral similarity is 0.7 or higher.

**Supplementary Figure 4.**
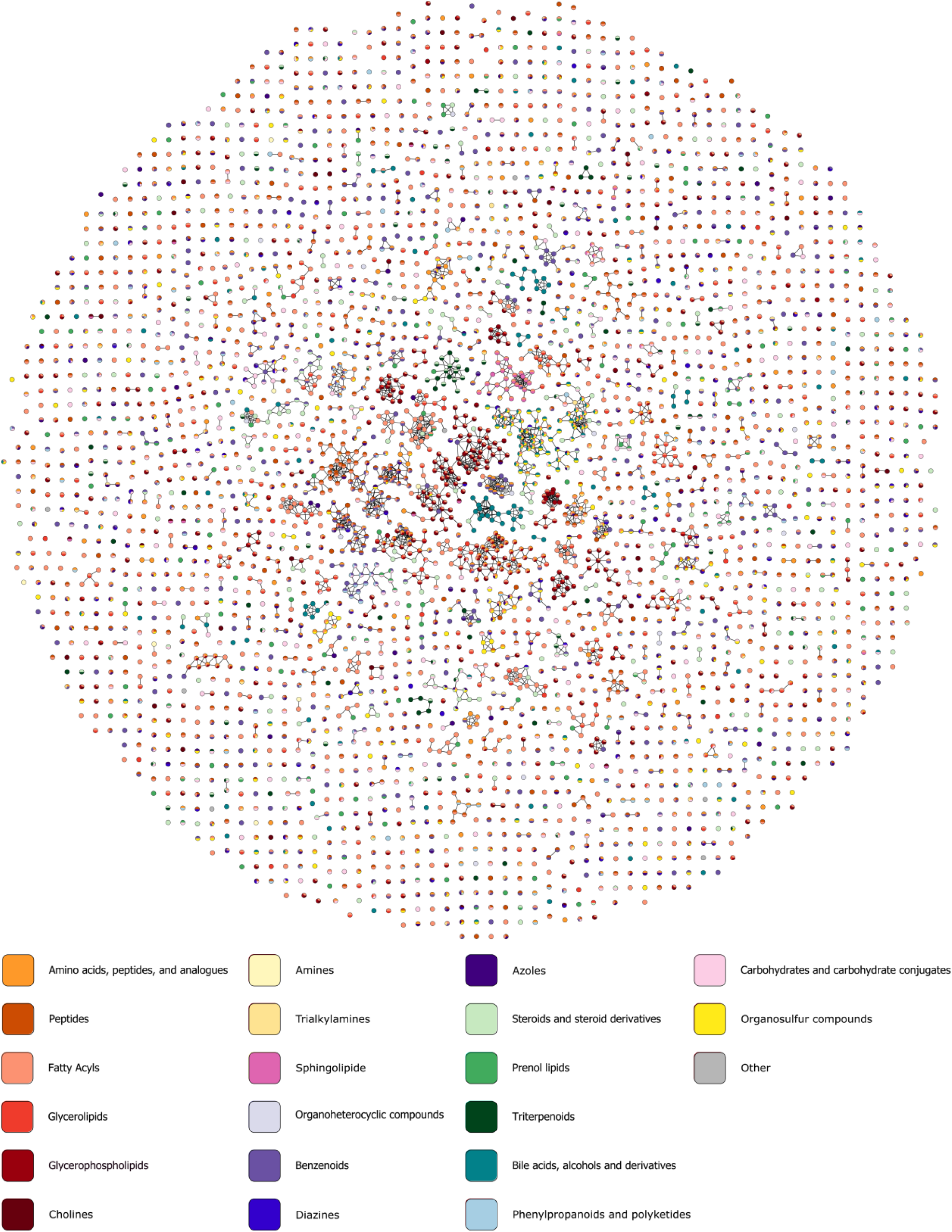
Molecular network and compound class annotations for the mice digestive system. Node colors indicate the compound class annotated by CANOPUS; compound classes are the same as in Supplementary Figure 3. Compounds belonging to multiple classes displayed as multicolored nodes. Nodes are connected by an edge if the spectral similarity is 0.7 or higher.

**Supplementary Figure 5.**
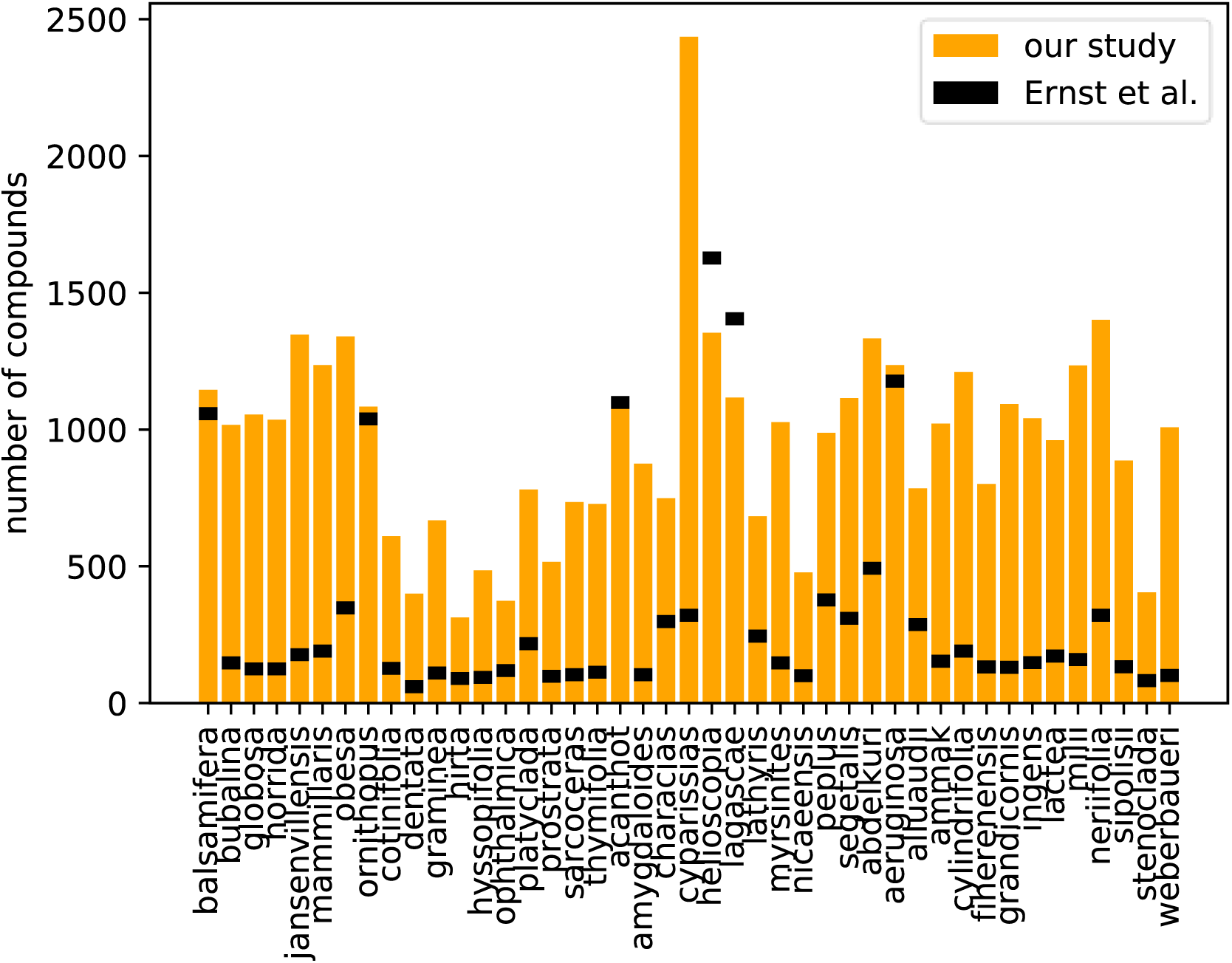
Number of compounds detected for each Euphorbia subgenus. Orange bars indicate the number of compounds detected here, black ticks indicate the number of compounds reported in the original study.

**Supplementary Figure 6.**
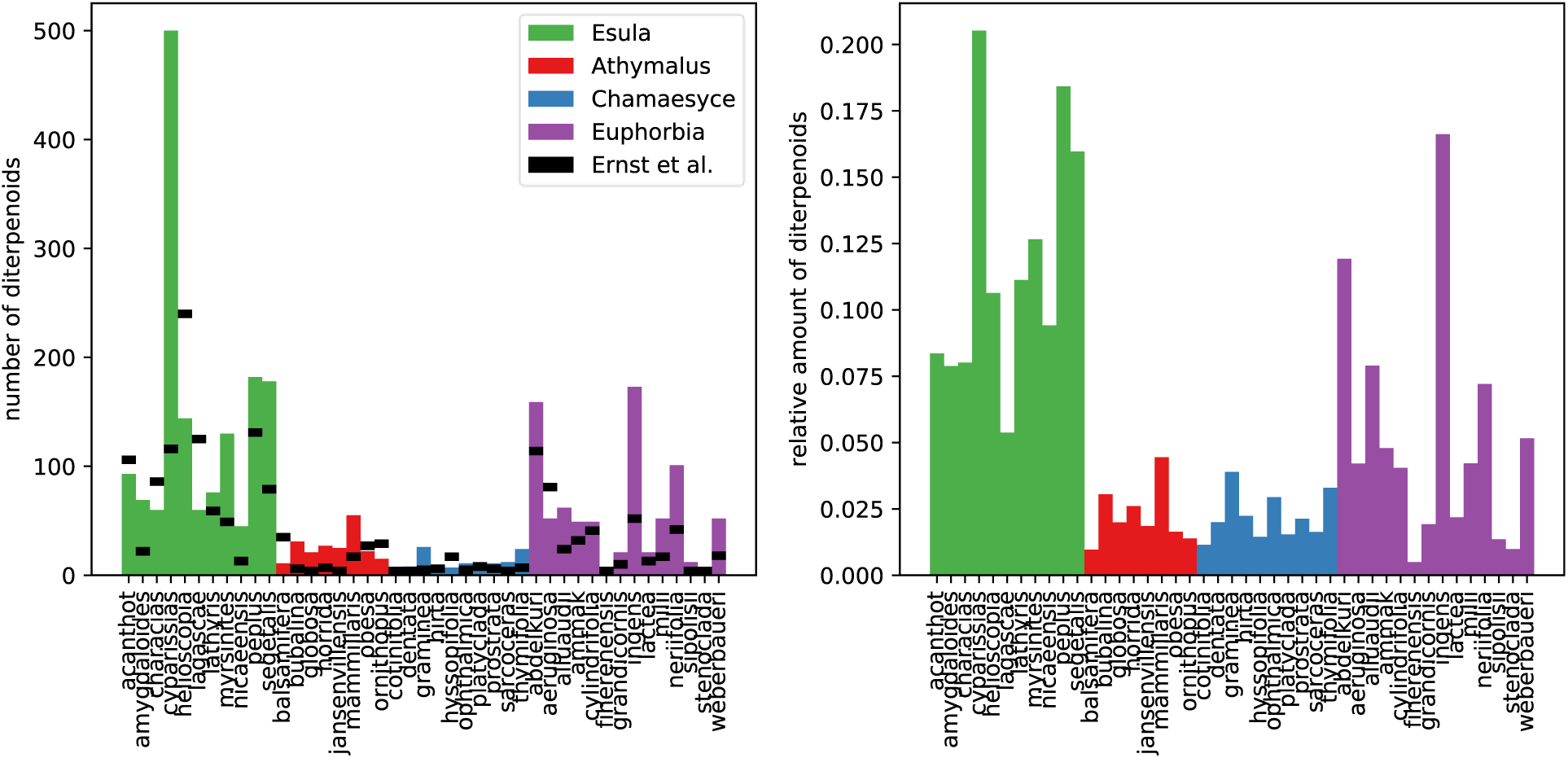
Number of compounds annotated as *diterpenoids* in different species of *Euphorbia*. Left: absolute number of compounds. Right: relative number of compounds, that is, number of *diterpenoids* divided by total number of compounds in each species. Black ticks in the left figure mark the reported number of *diterpenoids* in the original study by Ernst et al.

**Supplementary Figure 7.**
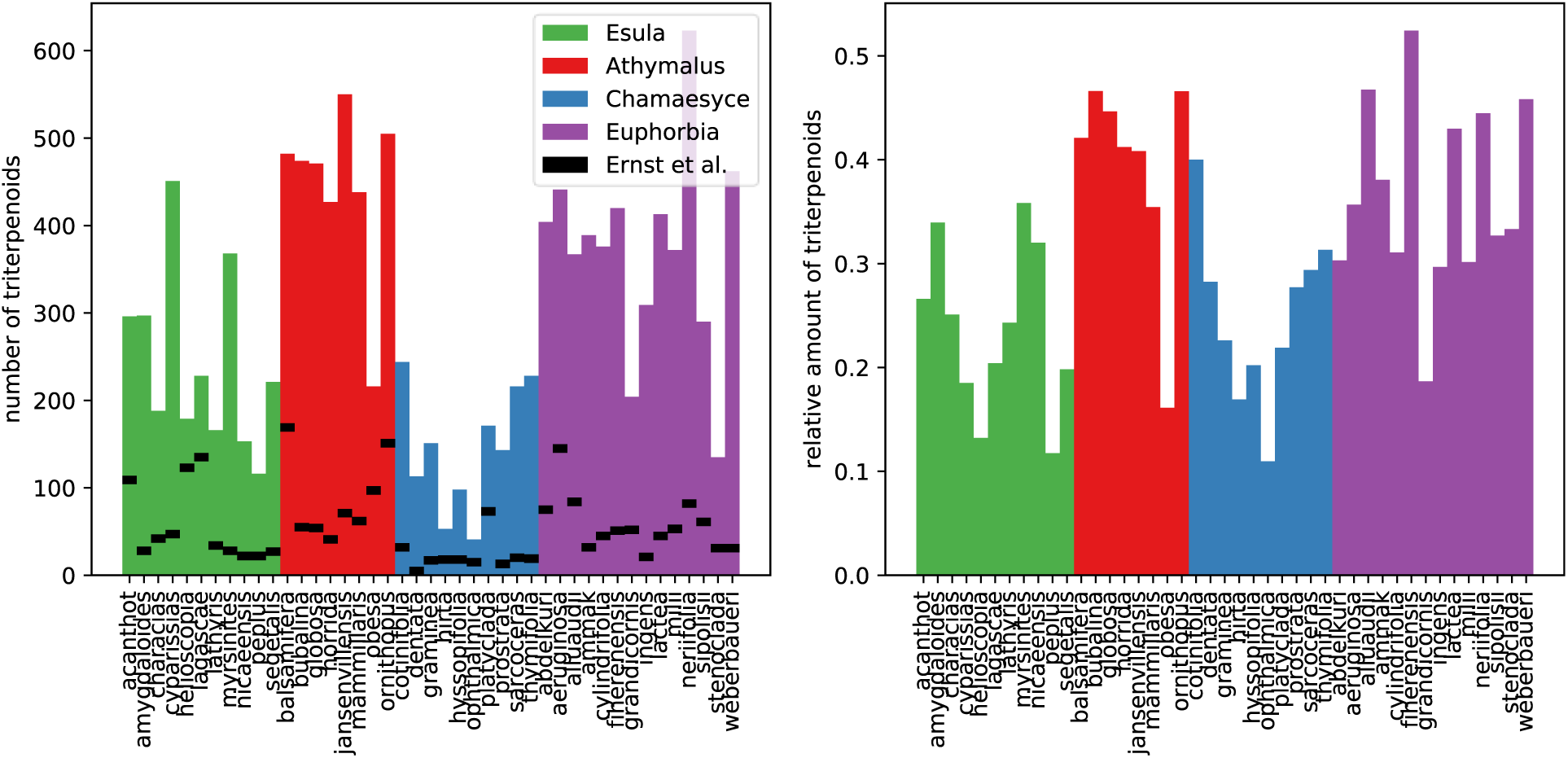
Number of compounds annotated as *triterpenoids* in different species of *Euphorbia*. Left: absolute number of compounds. Right: relative number of compounds, that is, number of *triterpenoids* divided by total number of compounds in each species. Black ticks in the left figure mark the reported number of *triterpenoids* in the original study by Ernst et al.

**Supplementary Figure 8.**
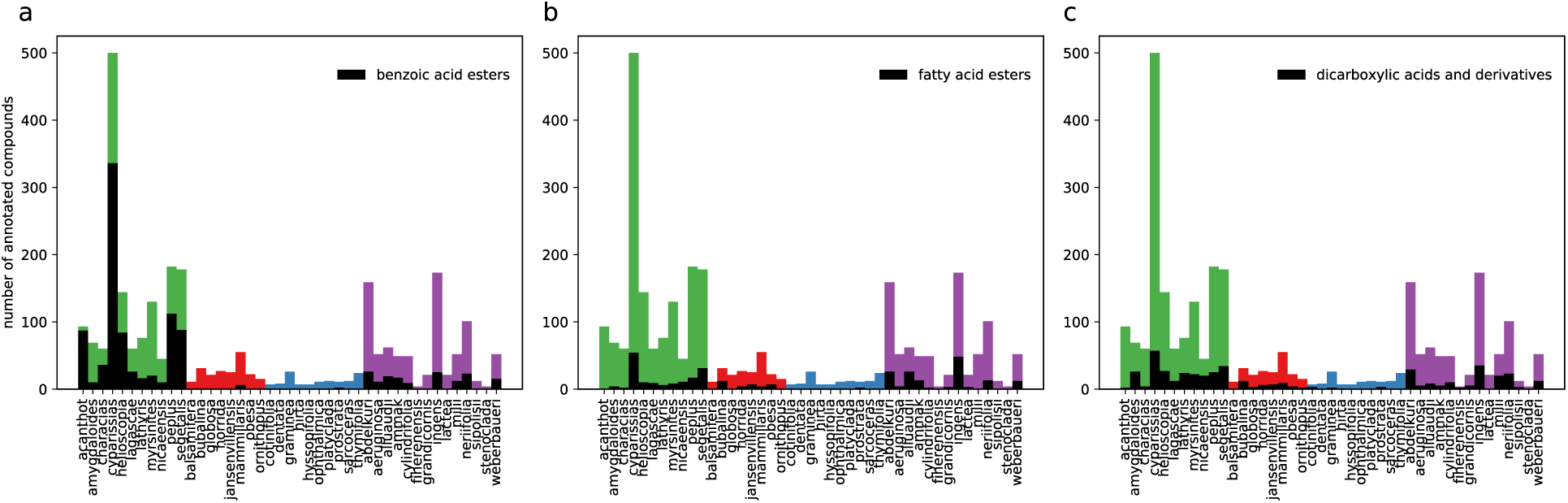
Number of *diterpenoids* in different species of *Euphorbia*. Black bars show the amount of *diterpenoids* that have a benzoic acid ester (a), fatty acid ester (b) or two carboxylic acids (c).

**Supplementary Table 2.** Features for the molecular formula input layer.

| Index | Description | Encoding |
| --- | --- | --- |
| 1-12 | Number of C, H, N, O, P, S, B, Cl, F, I, Br, Se | Integer |
| 13 | Exact mass | Real number |
| 14 | Ring-Double-Bond equivalent (RDBE) | Real number |
| 15 | RDBE is zero | Binary (-1, 1) |
| 16 | $\frac{RDBE}{(mass/100)^{2/3}}$ | Real number |
| 17 | Hetero-to-carbon ratio | Real number |
| 18 | Hetero-to-oxygen ratio | Real number |
| 19 | Hetero-without-oxygen-to-carbon ratio | Real number |
| 20 | Hydrogen-to-carbon ratio | Real number |
| 21 | Oxygen-to-carbon ratio | Real number |
| 22 | Nitrogen-to-carbon ratio | Real number |
| 23 | Molecular formula consists only of the elements C,H,N,O,P,S | Binary (-1, 1) |
| 24 | Molecular formula consists only of the elements C,H,N,O | Binary (-1, 1) |

This is file “evaluation_SVM_training_data.xlsx”

**Supplementary Material 1. Performance of CANOPUS and direct prediction for individual compound classes; evaluation on the SVM training dataset.** Performance results for all 1,270 compound classes from ClassyFire with at least 500 compounds with MS/MS data. We report accuracy, precision, recall, MCC and F1 score for each individual compound class. We also report evaluation results when removing *glycoflavonoids* and *bile acids* from the training data.

This is file “method_comparison_independent_data.xlsx”

**Supplementary Material 2. Performance of all evaluated methods for individual compound classes; evaluation on the independent dataset.** Performance results for all 1,270 compound classes from Supplementary Material 1. We report accuracy, precision, recall, MCC and F1 score for each individual compound class and each method.

This is file “plants_comparison.html”

**Supplementary Material 3. Interactive comparison of *Euphorbia* plants.** The user can select any two plant species to be compared; two sunburst plots then show the number of compounds annotated by CANOPUS for each compound class. Mouse-over allows to display details of a compound class, including the number and percentage of compounds that belong to this class, and the ClassyFire ontology and description of the class.

